# Intronic enhancers regulate the expression of genes involved in tissue-specific functions and homeostasis

**DOI:** 10.1101/2020.08.21.260836

**Authors:** Beatrice Borsari, Pablo Villegas-Mirón, Hafid Laayouni, Alba Segarra-Casas, Jaume Bertranpetit, Roderic Guigó, Sandra Acosta

## Abstract

Tissue function and homeostasis reflect the gene expression signature by which the combination of ubiquitous and tissue-specific genes contribute to the tissue maintenance and stimuli-responsive function. Enhancers are central to control this tissue-specific gene expression pattern. Here, we explore the correlation between the genomic location of enhancers and their role in tissue-specific gene expression. We found that enhancers showing tissue-specific activity are highly enriched in intronic regions and regulate the expression of genes involved in tissue-specific functions, while housekeeping genes are more often controlled by intergenic enhancers. Notably, an intergenic-to-intronic active enhancers continuum is observed in the transition from developmental to adult stages: the most differentiated tissues present higher rates of intronic enhancers, while the lowest rates are observed in embryonic stem cells. Altogether, our results suggest that the genomic location of active enhancers is key for the tissue-specific control of gene expression.

## Introduction

Multiple layers of molecular and cellular events tightly control the level, time and spatial distribution of expression of a particular gene. This wide range of mechanisms, known as gene regulation, defines tissue-specific gene expression signatures (Melé et al., 2015), which account for all the processes controlling the tissue function and maintenance, namely tissue homeostasis. Both the level and spatio-temporal pattern of expression of a gene are determined by a combination of regulatory elements (REs) controlling its transcriptional activation. Most genes contributing to tissue-specific expression signatures are actively transcribed in more than one tissue, but at different levels and with distinct patterns of expression in time and space, suggesting that the regulation of these genes is different across tissues. Nevertheless, approximately 10-20% of all genes are ubiquitously expressed (*housekeeping genes*), and they are involved in basic cell maintenance functions (Pervouchine et al., 2015; Zabidi et al., 2015; Eisenberg and Levanon, 2013).

*cis-*REs (CREs) are distributed across the whole genome, and changes in chromatin facilitate the transcriptional control over their target genes (Chen et al., 2019; Hawkins et al., 2010; Choukrallah et al., 2015). The activation of CREs depends on several epigenetic features, including combinations of different transcription factors’ binding sites, and it is positively correlated with the H3K27ac histone modification signal (Heinz et al., 2015; Heintzman et al., 2007). Epigenetic features in specific tissues may change throughout the life-span of individuals. During development, embryos undergo dramatic morphological and functional changes. These changes shape cell fate and identity as a result of tightly regulated transcriptional programs, which in turn are intimately associated with CREs’ activity and chromatin dynamics (Shlyueva et al., 2014; Bonev et al., 2017; Rand and Cedar, 2003; Gilbert et al., 2003).

Notably, key CREs known to regulate gene expression have been reported to locate in introns of their target genes (Ott et al., 2009; Kawase et al., 2011). However, it is unknown whether this is either a sporadic feature associated with certain types of genes - for instance long genes, such as HBB (*β*-globin) (Gillies et al., 1983) or CFTR (Ott et al., 2009) -, a common regulatory mechanism to most genes (Khandekar et al., 2007; Levine, 2010), or a pattern of biological significance. To delve into this question, we analyzed the genomic location of CREs across a panel of 87 adult and embryonic human cell types available from the Encyclopedia of DNA Elements (ENCODE) Project (Abascal et al., 2020). We found that highly shared CREs are mostly intergenic, while tissue-specific CREs tend to accumulate in introns. The prevalence of intronic CREs correlates with the level of specialization of the tissues, with the more differentiated ones presenting enrichment of intronic CREs. Moreover, intronic CREs target genes involved in tissue-specific functions and homeostasis, suggesting their implication in the functional specificity of tissues.

## Results

### Enhancer-like regulatory elements define tissue-specific signatures

We leveraged the cell type-agnostic registry of candidate *cis*-Regulatory Elements (cCREs) generated for the human genome (hg19) by the ENCODE Project. We focused on the set of 991,173 cCREs classified as Enhancer-Like Signatures (ELSs), defined as DNAse hypersensitive sites supported by the H3K27ac epigenetic signal, and assessed their presence-absence patterns across 60 adult cell type-specific catalogues (Table S1; see Methods). We first explored the data with multidimensional scaling (MDS), which uncovered tissue-specific presence-absence patterns (Fig. S1A). Indeed, the separation of samples driven by ELSs’ activity was comparable to the one obtained from the analysis of Genotype-Tissue Expression (GTEx) data (Melé et al., 2015), with blood and brain as the most diverging samples. This suggests a correlation between gene regulation mechanisms orchestrated by ELSs and tissue-specific gene expression patterns, which has been previously described (Pennacchio et al., 2007; Ernst et al., 2011).

Interestingly, we observed that the proportion of active ELSs located in intergenic regions was positively correlated with the number of samples in which ELSs were active (Spearman’s *ρ* = 0.55; *p* value = 6.2e-06; Fig. 1A), suggesting a functional role for the genomic location of ELSs. Thus, to untangle the relationship between genomic location and cell-type specificity of ELSs, we selected a subset of 25 samples that clustered into 5 main groups - iPSCs, fibro/myoblasts, muscle, blood and brain samples (Fig. 1B-C; Table S1, *Samples’ Cluster*) - according to their MDS proximity and consistently with their tissue of origin and function. This curated subset of samples allowed us to study enhancer activity in a tissue-specific manner, and compare it with regulatory mechanisms shared among tissues. Tissues represented by only one sample were not included in the subsequent analysis. Indeed, the fact that the *ad hoc* tissues’ functional clustering is supported by tissue-specific enhancer signatures suggests a direct link between ELSs’ activity and the regulation of tissue-specific functions. We defined *tissue-active* ELSs as those active in ≥ 80% of the samples within a given cluster (Table S2, **Tissue-active ELSs**; see Methods). As expected, in some cases we observed shared regulatory activity between tissues, in other words a fraction of ELSs active in a given cluster were also active in samples belonging to other clusters. For instance, approximately 1,700 blood-active ELSs were also active in all the seven brain samples (Fig. S1B). Because of this overlap, we defined sets of *tissue-specific* ELSs (Table S2, see Methods) as those active in ≥ 80% of the samples within the tissue cluster and in at most one sample outside the cluster. Due to their small size, for iPSC and fibro/myoblast clusters we considered as tissue-specific those ELSs active exclusively within their clusters (see Methods). The overlap of tissue-specific ELSs with samples from other clusters is depicted in Fig. 1D. The majority of brain- and blood-specific ELSs were active only within their tissue cluster (71.9% and 62.3%, respectively), while a considerable fraction (52.0%) of muscle-specific ELSs was shared with one sample from other clusters, mostly with fibro/myoblast samples (33.1%). This is consistent with the samples’ MDS proximity observed in Fig. 1B, suggesting a functional relevance of the genes regulated by shared ELSs. In addition, we identified a set of 208 ELSs active in all the 25 samples (Table S2, **Common ELSs**).

**Figure 1.**
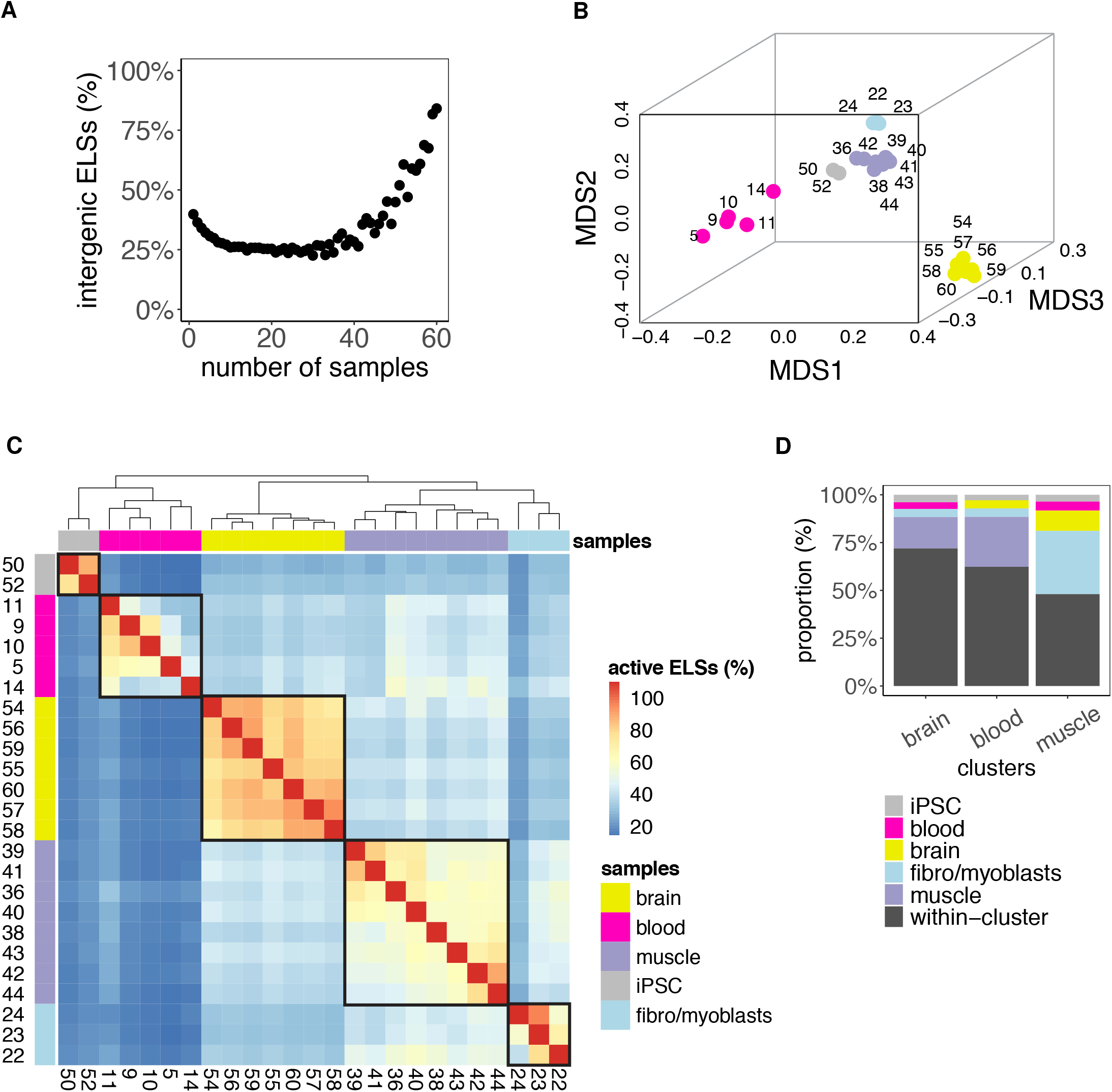
**A**: Highly-shared ELSs are more frequently located in intergenic regions. The scatter plot represents the proportion of intergenic ELSs active in increasing numbers of human adult samples (Spearman’s *ρ* = 0.55; *p* value = 6.2e-06). **B**: MDS distribution of human adult samples defined by ELSs’ activity. Analogous representation to Figure S1A for the subset of 25 selected adult human samples. **C**: Samples’ clustering defined by ELSs’ presence-absence patterns (clustering method: *complete*; clustering distance: *euclidean*). The heatmap represents the percentage of ELSs active in row *i* that are also active in column *j*. For this analysis we considered 268,214 of the 991,173 ELSs that were active in at least 2 of the 25 selected human adult samples. The correspondence between samples and numbers is reported in Table S1. **D**: Tissue-specific ELSs. The barplot represents the type of samples found within sets of brain-, blood- and muscle-specific ELSs. As described in Methods (section *Tissue-active, tissue-specific and common ELSs*), most of tissue-specific ELSs are only active in the samples of the corresponding cluster (“within-cluster”, *black*), but a few of them may be active in at most one outer sample (i.e. a sample that does not belong to the tissue cluster, *coloured*). IPSC- and fibro/myoblasts-specific ELSs are not represented, since we did not allow outer samples given their small cluster sizes (2 and 3, respectively; see Methods).

### The genomic location of regulatory elements correlates with their tissue-homeostatic functions

We next explored the genomic location of the sets of common and tissue-specific ELSs. While common ELSs were preferentially located in intergenic regions (63.4%, Fig. 2A), the majority of muscle- and brain-specific ELSs fell inside introns (71.6% and 74.0%, respectively; Fig. 2A). These significant differences in genomic distribution between tissue-specific and common regulatory elements (Table S3) are consistent with our initial observation of a high sharing rate of intergenic ELSs across samples (Fig. 1A). In contrast, the iPSC, fibro/myoblasts and blood clusters - which comprise undifferentiated, non-specialized or more heterogeneous cell types, respectively - showed a more even distribution of tissue-specific ELSs between intergenic and intronic regions (Fig. 2A). Overall, we observed a scarcity of exonic ELSs (Fig. 2A, Table S4).

**Figure 2.**
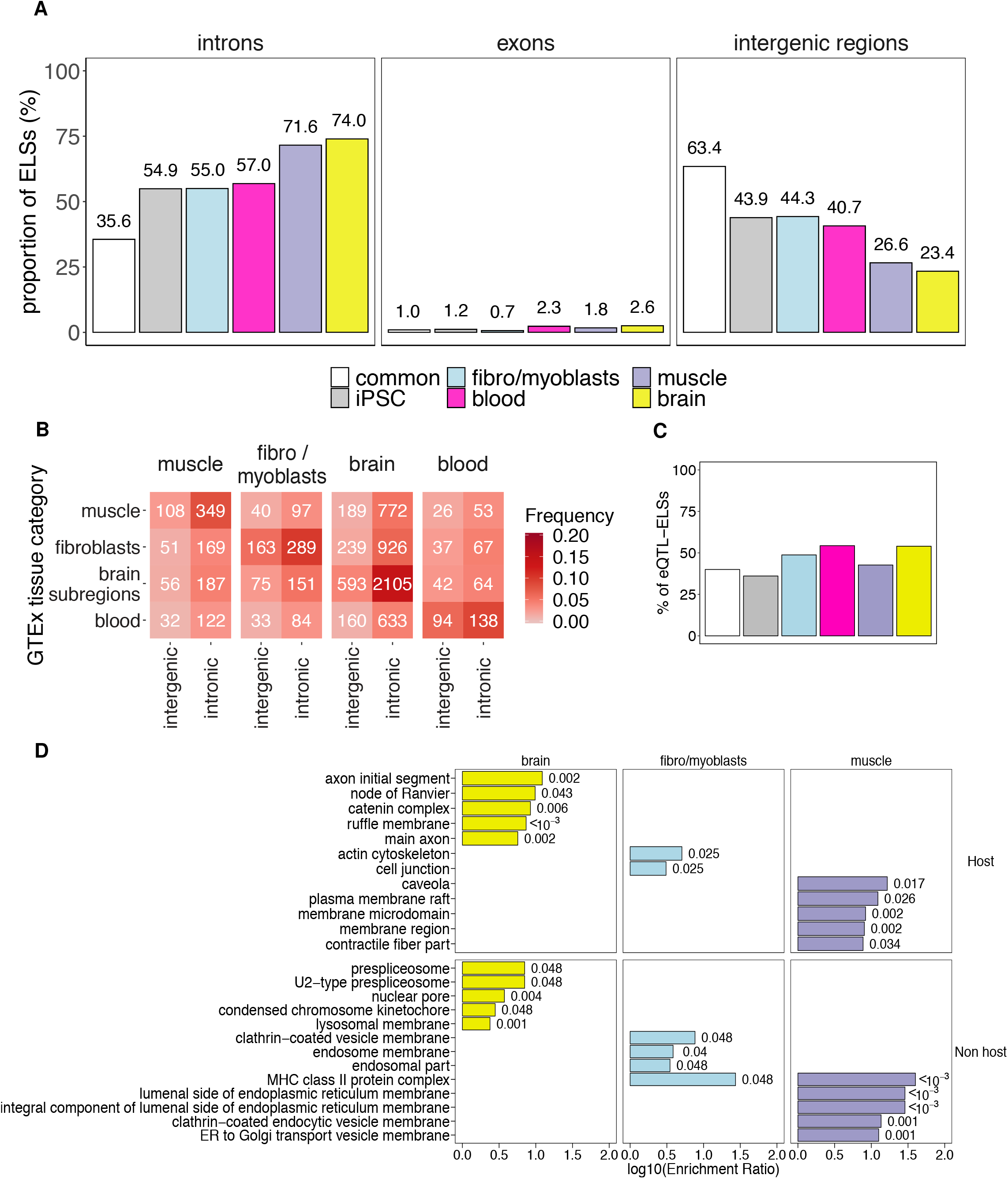
**A**: Proportions of common and tissue-specific ELSs identified in the 25 selected human adult samples that overlap intronic, exonic and intergenic regions. **B**: Number of intergenic and intronic muscle-, fibro/myobblasts-, brain- and blood-specific ELSs harboring eQTLs detected in Muscle, Fibroblasts, Brain subregions and Blood GTEx samples. Coloured cells represent the proportion of eQTL-ELSs over the total amount of tissue-specific ELSs within each group. **C**: Proportions of common and tissue-specific eQTL-ELSs targeting their host genes. These proportions were computed over the total amount of intronic eQTL-ELSs within each group. **D**: Top five enriched GO terms associated with the hosting and non-hosting eQTL-ELSs regulated genes. *P* value (FDR corrected) is reported for each enriched term.

Genes harboring tissue-specific ELSs may present distinctive features, including differences in intron length and density. To rule out any bias in our analyses, we compared these features between genes hosting common and tissue-specific ELSs. While the number of introns per hosting gene was comparable across groups (Kruskal-Wallis *p* value test = 0.98; Fig. S2A), we reported significant differences in the median intron length per gene (Kruskal-Wallis *p* value test < 2.2e-16; Fig. S2A). Moreover, we observed significant differences in the intronic ELSs’ density (Kruskal-Wallis *p* value test < 2.2e-16), with higher values for brain and muscle, suggesting that the enrichment of tissue-specific ELSs in intronic regions is not biased by the intron length (Fig. S2A).

We subsequently explored whether the genes harboring tissue-specific intronic ELSs perform functions associated with tissue homeostasis maintenance and response to stimuli. We performed a Gene Ontology (GO) enrichment analysis on the genes containing tissue-specific intronic ELSs. Indeed, the enrichment of terms associated with tissue-specific cellular components is consistent with the ELSs’ identity (Table S5). For instance, genes hosting brain-specific ELSs perform functions associated with synapses and axons, while in the case of muscle and blood we found significant terms related to sarcolemma, Z-disc and contractile fibers, and immunological synapses and cell membranes, respectively. Conversely, genes harboring common ELSs reported terms related to non-specific cell membrane composition (Table S5). Although this suggests an implication of intronic ELSs in tissue-specific functions, likely through tissue-specific gene regulation mechanisms, there is no proven association of intronic ELSs being direct regulators of their host genes.

To address this issue, we integrated our ELS analysis with the catalogue of expression Quantitative Trait Loci (eQTLs) provided by the Genotype-Tissue Expression (GTEx) Project (Aguet et al., 2017). Among the 35,275 common and tissue-specific ELSs, 5,941 overlap with a significantly associated eQTL-eGene pair, hereafter referred to as eQTL-ELSs. The proportion of eQTL-ELSs was similar among groups, with the exception of iPSC, which are not represented in the GTEx sampling collection (Fig. S2B). This allowed us to leverage the eQTL-ELSs pairs to explore the biological function of the genomic distribution of ELSs, focusing on eQTLs regulating gene expression in the four GTEx categories matching our samples’ clusters (fibroblasts, blood, muscle and brain subregions; see Methods). In line with the above-mentioned results, highly specialized tissues such as brain and muscle showed the highest proportion of intronic *vs* intergenic ELSs hosting eQTLs detected in the corresponding tissue: brain (2,105 (78%) *vs* 593 (22%)), muscle (349 (76%) *vs* 108 (24%)), fibro/myoblasts (289 (63%) *vs* 163 (37%)), blood (138 (59%) *vs* 94 (41%)) (Figure 2B). Conversely, common eQTL-ELSs were more frequently located in intergenic elements (5 (25%) *vs* 15 (75%)) (data not shown). Overall, these results indicate a potential functional role of the genomic distribution of ELSs in the regulation of tissue-specific gene expression. Still, although there is a clear trend of eQTL-ELSs’ specificity per tissue, many of these eQTLs are not exclusive to a single tissue. For this reason, we validated our observations with a GO enrichment analysis on the sets of genes associated with intronic and intergenic eQTL-ELSs. GO analysis on muscle- and brain-specific eQTL-ELSs showed a clear prevalence of tissue-specific homeostatic functions for those genes targeted by intronic eQTL-ELSs (for instance, muscle: carbohydrate and amino acid metabolism; brain: cell projection and organization). On the contrary, in the case of blood we found significantly enriched GO terms only for genes targeted by intergenic eQTL-ELSs (Table S6). This might be due to the fact that blood comprises different cell types and can be considered a more heterogeneous tissue. Overall, these results suggest that intronic eQTL-ELSs are involved in the regulation of genes controlling tissue-specific functions and tissue homeostasis.

Next, we wanted to understand the relationship between the intronic ELSs and their harboring genes. Of note, the proportion of intronic eQTL-ELSs targeting their host genes was comparable among groups of samples, but always below the 54.3% (Fig. 2C). Most interestingly, eQTL-ELSs regulating the expression of the host gene are associated with tissue-specific functions, with genes involved in axonal components for the brain (e.g. NRCAM), actin cytoskeleton for fibroblasts (e.g. FMN1) or contractility-related terms for muscle (e.g. SYNM). However, those targeting the expression of non-hosting genes are involved in homeostatic functions not directly associated with the tissue function. For instance, the brain presents significant terms related to the splicing proteins (e.g. SF3A1, SF3B1), a widely extended process in the brain and responsible of the fine tuning of several brain functions (Vuong et al., 2016) (Fig. 2D). Overall, this suggests that other mechanistic strategies may account for the intronic preference of regulatory elements in highly specialized tissues.

### The enrichment of transcription factor binding sites in tissue-specific ELSs is independent of their genomic location

The activation of ELSs is a dynamic process depending, mainly, on its accessible chromatin to be bound by transcription factors (TFs). Thus, tissue-specific gene expression programs may be controlled by the underlying signature of TFs-ELSs pairing (Schmitt et al., 2016). We next wondered whether the specific distribution of ELSs, i.e. intronic *vs* intergenic, was associated with a different transcription factor binding site (TFBS) signature that could account for their tissue specificity. For this purpose we explored, using the software HOMER (Heinz et al., 2010), TFBSs differences between intronic and intergenic ELSs that were either common or specific to a given tissue. We observed a high sharing rate of TFBSs between intronic and intergenic ELSs, suggesting that there is a strong prevalence of certain transcriptional programs in each tissue independently of the genomic location of ELSs. Notably, there are no enriched TFBSs in common ELSs, either intronic or intergenic (Fig. 3A and Table S7). Amongst the TFBSs enriched in the tissue-specific intronic and intergenic ELSs, there are some that are well known to control tissue-specific homeostatic events, such as FLI1 and RUNX in blood controlling adult endothelial hemogenesis (Lis et al., 2017), and POU6F1 (Brn5), SOX4 and SOX8 in brain controlling the adult neural plasticity (McClard et al., 2018). POU5F1 (Oct4) is required for iPSC reprogramming, and MEIS1 in muscle is key for cardiomyogenesis (Dupays et al., 2015). Although a great number of the TFs identified in our analysis are known for shaping the functions of certain tissues, the vast majority of these TFs are ubiquitously or widely expressed in several tissues (Fig. 3B), suggesting that the tissue-specificity of gene regulation does not arise from the transcription factor’s potential to bind an ELS, but most likely from the genomic localization of the ELSs.

**Figure 3.**
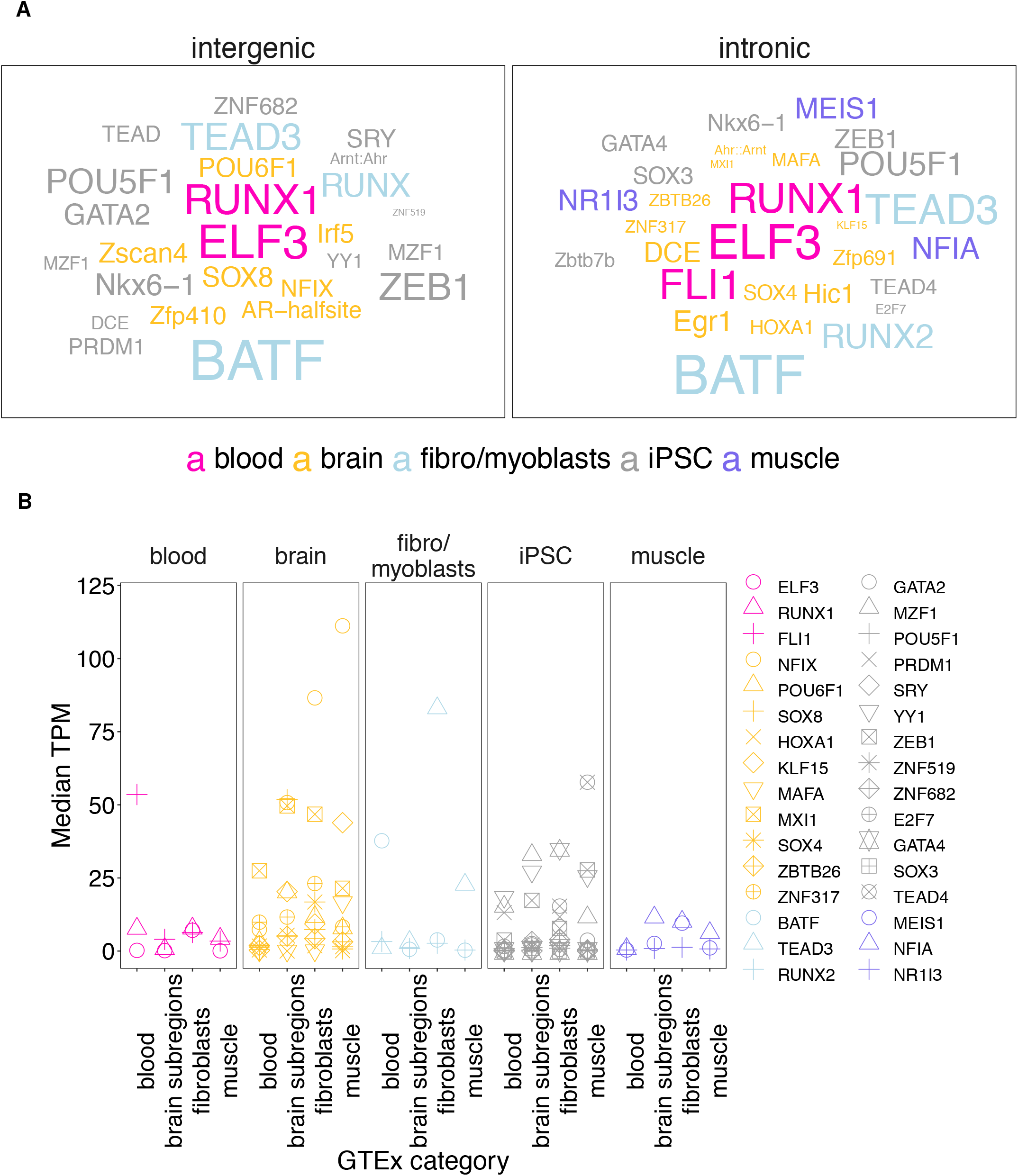
**A**: Word cloud reporting the TFBSs significantly enriched in intronic and intergenic tissue-specific ELSs. No significant TFBSs were found in common ELSs. The size of the word represents the significance of TFBSs enrichment. **B**: Median expression, in the four matching GTEx tissues categories, of the TFs associated with significantly enriched TFBSs in each cluster.

### The genomic location of developmental ELSs is not associated with tissue specificity

Tissue-specific homeostatic features vary dramatically among different adult tissues. For instance, blood comprises a number of cell types characterized by heterogeneous functions and high turnover. On the other hand, muscles are formed by fewer cell types, mainly dedicated to the same function and with limited cell division capacity. The maintenance of tissue homeostasis is ensured by quiescent adult stem cells with features similar to their developmental native lineage (Rué and Martinez Arias, 2015; Biteau et al., 2011). During development, tissues mature to fully reach their functional capacity in adulthood. Still, whether the regulatory features of a given tissue are reminiscent of their developmental lineage remains largely unknown. For this reason, we assessed the activity of the 991,173 cell type-agnostic ELSs across 27 embryonic samples (Table S8). The correlation between the percentage of intergenic ELSs and the number of samples in which ELSs are active was lower compared to adult samples (Spearman’s *ρ* = 0.38; *p* value = 0.054; Fig. 4A). MDS analysis highlighted three main groups of embryonic samples: stem cells (ESC), neural progenitors, and a heterogeneous group of more differentiated cell types (Fig. 4B; Table S8, Samples’ Group). The three groups of samples were associated with 3,112, 784 and 1,166 specific ELSs, respectively (Table S9). Although the majority of these ELSs were active only within the corresponding cluster, we reported that 26.2% of the neural progenitors-specific ELSs were also active in one ESC sample (Fig. S3A). On the contrary, we identified only 94 ELSs common to all embryonic samples (Table S9). The proportion of specific intronic ELSs was higher for neural progenitors and differentiated tissues (60.3% and 60.6%, respectively; Fig. 4C) compared to ESC-specific (50.9%) and common (38.3%) ELSs, but lower with respect to clusters of adult muscle and brain samples (71.6% and 74.0%, respectively, Fig. 2A). As in the case of adult samples, we observed a scarcity of exonic ELSs (Fig. 4C, Table S11), while we could not find significant associations between the frequency of group-specific intronic ELSs and features of intron length and density (Figs. S3B). On the other hand, the density of ELSs per introns (Fig. S3B) was similar to the one observed in adult samples (Fig. S2A).

**Figure 4.**
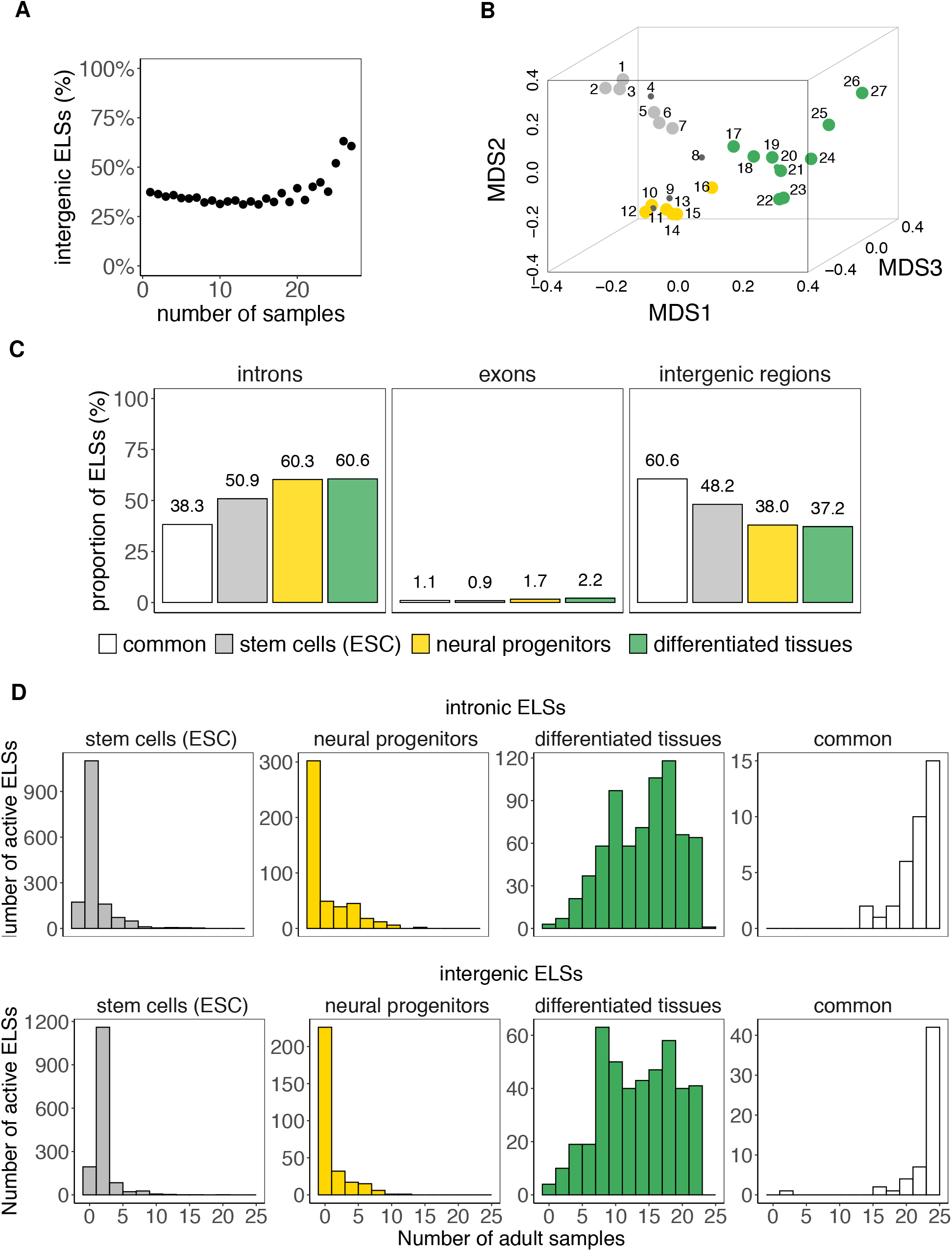
**A**: Scatter plot representing the percentage of intergenic ELSs active in increasing numbers of human embryonic samples (Spearman’s *ρ* = 0.38; *p* value = 0.054). The degree of correlation between ELSs’ sharing and the percentage of intergenic ELSs is lower compared to the one observed for adult samples. **B**: MDS representation of the dissimilarities between the 27 human embryonic samples according to the pattern of activity of ELS-cREs (analogous to Fig. 1B). The correspondence between samples and numbers is reported in Table S8. The MDS highlights 3 main groups of embryonic samples. **C**: Proportions of common and group-specific ELSs identified in embryonic samples that overlap intronic, exonic and intergenic regions. **D**: Rate of sharing of intronic (upper panel) and intergenic (lower panel) ELSs between embryonic and adult samples. The histogram represents the number of selected adult samples (n = 25) in which embryonic ELSs are active.

When studying the genes harboring developmental group-specific intronic ELSs, we observed that they are enriched in functions consistent with the corresponding adult tissue (Table S12). For instance, the ones hosting neural progenitors-specific ELSs are enriched in neural development-related terms, such as axonogenesis and dendritic spine organization. Notably, genes harboring developmental common ELSs are enriched in protein complexes like nBAF and SWI/SNF, known developmental chromatin remodelers (Alver et al., 2017).

Lastly, in an attempt to define the amount of regulatory activity shared by embryonic and adult samples as an indicator of the reminiscent embryonic function in adult tissue homeostasis, we computed, for specific and common embryonic ELSs, the number of adult tissues in which they were found active. As expected, whereas ELSs specific to stem cells and neural progenitors were active in a limited set of adult samples, embryonic differentiated tissues reported a higher degree of shared regulatory activity with adult cell types. Moreover, ELSs active in all embryonic samples (common) were also active in the majority of adult samples (Fig. 4D). Overall, these results show that the genomic location of ELSs is dynamic throughout development, and shifts towards intronic localization during tissue maturation.

## Discussion

In this study, we show the central role of intronic Enhancer-Like Signatures (ELSs) in the control of tissue-specific expression signatures. Tissue-specific homeostasis is a dynamic process encompassing the coordinated expression in time and space of a wealth of genes, mainly controlled by active ELSs. ENCODE data suggests that about half of these ELSs are intergenic, and 38% are intronic (ENCODE SCREEN Portal: https://screen-v10.wenglab.org/, section “About”). The enrichment in intronic ELSs in the most specialized tissues observed in our study, independently of the sequence - in terms of transcription factor binding sites - suggests an important role of the genomic location of ELSs. Since Heitz described in 1928 (Heitz, 1928) euchromatin as chromosomal regions enriched in genes, and heterochromatin as inactive or passive chromatin regions, this dual definition has been shaped throughout the years but it still remains vastly correct (De Laat and Duboule, 2013; DeMare et al., 2013; Ernst and Kellis, 2010). Intergenic regions are often transcriptionally and regulatory silenced, and notably they are more frequent in adult than embryonic tissues (Heinz et al., 2015). A similar correlation is observed in our data, since embryonic ELSs are not so frequently found in intronic elements as in adults, suggesting that the maturation and tissue commitment correlated with the ELS distribution across the whole genome. One could hypothesize that the enriched presence of intronic ELSs in specialized tissues is advantageous for the control of the gene expression signature of a particular tissue, for instance granting ELSs accessibility in open DNA regions (genes) and avoiding leaky activity of ELSs. Introns have been long observed as gene expression regulators throughout different mechanisms (Rose, 2019; Chorev and Carmel, 2012; Shaul, 2017). Introns regulatory potential has been longly associated with the regulation of the host gene’s expression in several different ways, often related to alternative splicing, intron retention (Jacob and Smith, 2017), non-sense mediated decay (Lewis et al., 2003), and even with the control of transcription initiation via recruitment of RNA Polymerase II (Bieberstein et al., 2012). However, here we found that about half of the eQTL-ELSs located in introns do not regulate the expression of the host gene. This is important regulatory information since it disentangles the presence of intronic ELSs from the regulation of the host gene, opening new opportunities to identify the regulatory mechanisms controlling tissue-specific gene expression. Overall, our results suggest that the genomic distribution of tissue-specific active ELSs is not stochastic and mainly overlaps with intronic elements. The opposite happens to active ELSs common to all tissues. These results suggest that introns play a role in the regulation of gene expression in a tissue-specific manner.

## Methods

### The ENCODE registry of candidate *cis*-Regulatory Elements

The cell type-agnostic registry of human candidate *cis*-Regulatory Elements (cCREs) available from the ENCODE portal corresponds to a subset of 1,310,152 representative DNase hypersensitivity sites (rDHSs) in the human genome with epigenetic activity further supported by histone modification (H3K4me3 and H3K27ac) or CTCF-binding data (https://screen-v10.wenglab.org/; section “About”). It comprises 991,173 Enhancer-Like Signatures (ELS), 254,880 Promoter-Like Signatures (PLS), and 64,099 CTCF-only Signatures. In addition, cell type-specific catalogues are provided for those cell types with available DNase and ChIP-seq ENCODE data.

### Selection of cCREs with enhancer-like signature (ELS) across human samples

We downloaded the set of 1,310,152 cell type-agnostic cCREs for human assembly 19 (hg19) from the ENCODE SCREEN webpage (https://screen-v10.wenglab.org/; file ID: ENCFF788SJC). From the ENCODE portal (https://www.encodeproject.org/matrix/?type=Annotation&encyclopedia_version=ENCODE+v4&annotation_type=candidate+Cis-Regulatory+Elements&assembly=hg19), we retrieved cell type-specific registries of cCREs for 60 adult and 27 embryonic human samples with available DNase data and ChIP-seq H3K4me3 and H3K27ac data. The ENCODE File Identifiers for the adult and embryonic datasets are reported in Table S1 and S8, respectively. We focused on the 991,173 cell type-agnostic cCREs with ELS activity, and generated a binary table in which we assessed, for a given cCRE, the presence/absence of ELS activity annotation (column 9 = “255, 205, 0”) in each of the 60 adult and 27 embryonic samples. A binary distance matrix between all pairs of adult samples was used to perform multidimensional scaling (MDS) in three dimensions. This resulted in the selection of 25 adult samples. The same procedure was applied, independently, to the embryonic samples. In this case, IMR-90, mesendoderm, mesodermal cell, endodermal cell and ectodermal cell samples were not included in subsequent analyses.

### Intersection of ELSs with genes, introns, exons and intergenic regions

Genes, exons and introns’ coordinates were obtained from GENCODE v19 annotation (https://www.gencodegenes.org/human/release_19.html). The overlap between ELSs and genes, exons and introns was computed using BEDTools intersectBed v2.27.1 (Quinlan and Hall, 2010). The proportions of ELSs overlapping intronic segments (Figs. 2A, 4C) also include a limited set of ELSs overlapping both intronic and exonic regions (common adult ELSs: 2.4%; iPSC-specific ELSs: 3.1%; fibro/myoblasts-specific ELSs: 4.5%; blood-specific ELSs: 5.6%; muscle-specific ELSs: 4.4%; brain-specific ELSs: 7.4%; common embryonic ELSs: 7.4%; differentiated tissues-specific ELSs: 5.1%; neural progenitors-specific ELSs: 5.0%; ESC-specific ELSs: 3.2%). On the other hand, we defined as exonic ELSs those intersecting exclusively exonic regions (Figs. 2A, 4C). The overlap of ELSs with intergenic regions was obtained by intersecting the former with the genes’ coordinates using the BEDTools intersectBed option -v.

### Tissue-active, tissue-specific and common ELSs

Tissue-active ELSs are ELSs active (see Methods section *Selection of cCREs with enhancer-like signature (ELS) across human samples*) in ≥ 80% of the samples within a given group of samples (blood = 4/5; muscle = 6/8; brain = 6/7; stem cells = 5/6; neural progenitors = 5/6; differentiated tissues = 8/10). Because of the small sample size, we required iPSC- and fibro/myoblasts ELSs to be active in 100% of the samples (2/2; 3/3). Tissue-specific ELSs are tissue-active ELSs that are active in 0 (iPSC, fibro/myoblasts) or at most 1 (all other groups) outer samples (i.e. samples outside the considered group). Common adult and embryonic ELSs are ELSs active in 100% of the samples (25/25 and 22/22, respectively). To rule out indirect effects of ELS activity related to promoter regions, we discarded common and tissue-specific ELSs overlapping any annotated Transcription Start Site (TSS, ± 2Kb) in GENCODE v19.

### Assessing enhancer regulatory activity

ELSs were annotated by using the GTEx v7 (Aguet et al., 2017)) significant variant-gene pairs from 46 different tissues (number of samples with genotype ≥ 70). Only single-tissue eQTL-eGene associations with a qval ≤ 0.05 were used. Similar GTEx tissues were grouped in unique categories in order to consider the most complete catalogue of eQTL-eGene pairs per group of samples. These categories were named as follows: fibroblasts (Skin Not Sun Exposed Suprapubic, Cells Transformed Fibroblasts), blood (Whole Blood, Spleen), muscle (Skeletal Muscle), brain subregions (all brain subregions, Pituitary Gland, Nerve Tibial), cardiovascular (Heart Atrial Appendage, Heart Left Ventricle, Artery Aorta, Artery Coronary, Artery Tibial), digestive (Liver, Pancreas, Small Intestine Terminal Ileum, Stomach, Colon Sigmoid, Colon Transverse, Esophagus Gastroesophageal Junction, Esophagus Mucosa, Esophagus Muscularis, Adipose Subcutaneous, Adipose Visceral Omentum), gland (Adrenal Gland, Thyroid, Minor Salivary Gland), breast (Breast Mammary Tissue), lung (Lung), sexual (Ovary, Prostate, Testis, Uterus, Vagina). Bedtools (Quinlan and Hall, 2010) was used to intersect the tissue-specific ELSs’ coordinates with the *cis*-eQTLs’ positions in the considered genomic locations (intronic and intergenic). We kept all eQTL-eGene pairs that were found significantly associated with the matching eQTL-ELS’s tissue category (brain, blood, muscle and fibro/myoblasts). In the case of iPSC-specific and common ELSs, we considered those eQTL-eGene pairs that were significantly reported in all the tissues. The resulting intersected ELSs were considered as being responsible for the regulation of the associated eGene. The functional enrichment of the ELSs’ target genes was performed by the online utility WebGestalt (Liao et al., 2019).

### *cis*-Regulatory Elements and Transcription Factor Binding Sites

Transcription factor binding sites (TFBSs) were predicted by using the motif discovery software HOMER (Heinz et al., 2010) This program performs a differential motif discovery by taking two sets of genomic regions (findMotifGenome.pl script) and identifying the motifs that are enriched in one set of sequences relative to a background list of regions. We analysed the tissue-specific ELSs’ binding motifs by considering the ELS regions from all the other tissues as background. We searched for 6-mer and 7-mer length motifs as a way to focus on enriched core motif sequences and avoid redundancy from longer motifs with similar functions. A hypergeometric test and FDR correction were applied for the motif enrichment. Only significantly enriched motifs were considered in the subsequent analysis. The word size in Figure 3A is proportional to the significance of the enrichment, it is calculated as the difference of sequence frequencies where the TFBS is found in the target and background lists of regions. The functionality of the predicted TFBSs was assessed by analysing the tissue-specific expression of the transcription factors that bind to them. GTEx expression data (v7) was analysed for those transcription factors whose TFBSs were reported as significant by HOMER in all tissues and genomic locations.

## Data access

All ENCODE data used in this study is publicly available on the ENCODE portal (https://www.encodeproject.org/). GTEx gene expression and eQTL data is available on the GTEx portal (https://www.gtexportal.org).

## Acknowledgments

B.B. is supported by the fellowship 2017FI B00722 from the Secretaría d’Universitats i Recerca del Departament d’Empresa i Coneixement (Generalitat de Catalunya) and the European Social Fund (ESF). P.V-M. is supported by an FPI PhD fellowship (FPI-BES-2016-077706) part of the “Unidad de Excelencia María de Maeztu” funded by the MINECO (ref: MDM-2014-0370). S.A. is supported by a fellowship from the Secretaría d’Universitats i Recerca del Departament d’Empresa i Coneixement (Generalitat de Catalunya) (BP-2017-00176). We thank the ENCODE and GTEx Consortia for data production. We thank Diego Garrido-Martín (R. Guigó Lab) for valuable statistical advice.

## Author Contributions

S.A., B.B, and P.V-M. designed the study, analyzed the data and wrote the manuscript with feedback from all the authors. H.L. and A.S-C. analyzed the data.

## Competing interest statement

The authors declare no competing interests.

## References

Abascal, F., Acosta, R., Addleman, N. J., Adrian, J., Afzal, V., Aken, B., Akiyama, J. A., Jammal, O. A., Amrhein, H., Anderson, S. M., et al. (2020). Expanded encyclopaedias of DNA elements in the human and mouse genomes. Nature, 583(7818):699–710.

Aguet, F., Brown, A. A., Castel, S. E., Davis, J. R., He, Y., Jo, B., Mohammadi, P., Park, Y. S., Parsana, P., Segrè, A. V., et al. (2017). Genetic effects on gene expression across human tissues. Nature, 550(7675):204–213.

Alver, B. H., Kim, K. H., Lu, P., Wang, X., Manchester, H. E., Wang, W., Haswell, J. R., Park, P. J., and Roberts, C. W. M. (2017). The SWI/SNF chromatin remodelling complex is required for maintenance of lineage specific enhancers. Nature Communications, 8(1):1–10.

Bieberstein, N. I., Oesterreich, F. C., Straube, K., and Neugebauer, K. M. (2012). First exon length controls active chromatin signatures and transcription. Cell Reports, 2(1):62–68.

Biteau, B., Hochmuth, C. E., and Jasper, H. (2011). Maintaining tissue homeostasis: Dynamic control of somatic stem cell activity.

Bonev, B., Mendelson Cohen, N., Szabo, Q., Fritsch, L., Papadopoulos, G. L., Lubling, Y., Xu, X., Lv, X., Hugnot, J. P., Tanay, A., et al. (2017). Multiscale 3D Genome Rewiring during Mouse Neural Development. Cell, 171(3):557–572.

Chen, C., Yu, W., Tober, J., Blobel, G. A., Speck, N. A., and Correspondence, K. T. (2019). Spatial Genome Re-organization between Fetal and Adult Hematopoietic Stem Cells.

Chorev, M. and Carmel, L. (2012). The function of introns. Frontiers in Genetics, 3(APR):55.

Choukrallah, M. A., Song, S., Rolink, A. G., Burger, L., and Matthias, P. (2015). Enhancer repertoires are reshaped independently of early priming and heterochromatin dynamics during B cell differentiation. Nature Communications, 6(1):1–11.

De Laat, W. and Duboule, D. (2013). Topology of mammalian developmental enhancers and their regulatory landscapes.

DeMare, L. E., Leng, J., Cotney, J., Reilly, S. K., Yin, J., Sarro, R., and Noonan, J. P. (2013). The genomic landscape of cohesin-Associated chromatin interactions. Genome Research, 23(8):1224–1234.

Dupays, L., Shang, C., Wilson, R., Kotecha, S., Wood, S., Towers, N., and Mohun, T. (2015). Sequential Binding of MEIS1 and NKX2-5 on the Popdc2 Gene: A Mechanism for Spatiotemporal Regulation of Enhancers during Cardiogenesis. Cell Reports, 13(1):183–195.

Eisenberg, E. and Levanon, E. Y. (2013). Human housekeeping genes, revisited.

Ernst, J. and Kellis, M. (2010). Discovery and characterization of chromatin states for systematic annotation of the human genome. Nature Biotechnology, 28(8):817–825.

Ernst, J., Kheradpour, P., Mikkelsen, T. S., Shoresh, N., Ward, L. D., Epstein, C. B., Zhang, X., Wang, L., Issner, R., Coyne, M., et al. (2011). Mapping and analysis of chromatin state dynamics in nine human cell types. Nature, 473(7345):43–49.

Gilbert, N., Boyle, S., Sutherland, H., Heras, J. d.L., Allan, J., Jenuwein, T., and Bickmore, W. A. (2003). Formation of facultative heterochromatin in the absence of HP1. The EMBO Journal, 22(20):5540–5550.

Gillies, S. D., Morrison, S. L., Oi, V. T., and Tonegawa, S. (1983). A Tissue-specific Transcription Enhancer Element Is Located in the Major lntron of a Rearranged lmmunoglobulin Heavy Chain Gene. Technical report.

Hawkins, R. D., Hon, G. C., Lee, L. K., Ngo, Q., Lister, R., Pelizzola, M., Edsall, L. E., Kuan, S., Luu, Y., Klugman, S., et al. (2010). Distinct epigenomic landscapes of pluripotent and lineage-committed human cells. Cell Stem Cell, 6(5):479–491.

Heintzman, N. D., Stuart, R. K., Hon, G., Fu, Y., Ching, C. W., Hawkins, R. D., Barrera, L. O., Van Calcar, S., Qu, C., Ching, K. A., et al. (2007). Distinct and predictive chromatin signatures of transcriptional promoters and enhancers in the human genome. Nature Genetics, 39(3):311–318.

Heinz, S., Benner, C., Spann, N., Bertolino, E., Lin, Y. C., Laslo, P., Cheng, J. X., Murre, C., Singh, H., and Glass, C. K. (2010). Simple combinations of lineage-determining transcription factors prime cis-regulatory elements required for macrophage and B cell identities. Molecular cell, 38(4):576–589.

Heinz, S., Romanoski, C. E., Benner, C., and Glass, C. K. (2015). The selection and function of cell type-specific enhancers.

Heitz, E. (1928). Das Heterochromatin der Moose. Jahrbücher für wissenschaftliche Botanik, 69.

Jacob, A. G. and Smith, C. W. J. (2017). Intron retention as a component of regulated gene expression programs.

Kawase, S., Imai, T., Miyauchi-Hara, C., Yaguchi, K., Nishimoto, Y., Fukami, S. I., Matsuzaki, Y., Miyawaki, A., Itohara, S., and Okano, H. (2011). Identification of a novel intronic enhancer responsible for the transcriptional regulation of musashi1 in neural stem/progenitor cells. Molecular Brain, 4(1):14.

Khandekar, M., Brandt, W., Zhou, Y., Dagenais, S., Glover, T. W., Suzuki, N., Shimizu, R., Yamamoto, M., Lim, K. C., and Engel, J. D. (2007). A Gata2 intronic enhancer confers its pan-endothelia-specific regulation. Development, 134(9):1703–1712.

Levine, M. (2010). Transcriptional enhancers in animal development and evolution.

Lewis, B. P., Green, R. E., and Brenner, S. E. (2003). Evidence for the widespread coupling of alternative splicing and nonsense-mediated mRNA decay in humans. Proceedings of the National Academy of Sciences of the United States of America, 100(1):189–192.

Liao, Y., Wang, J., Jaehnig, E. J., Shi, Z., and Zhang, B. (2019). WebGestalt 2019: gene set analysis toolkit with revamped UIs and APIs. Nucleic Acids Research, 47(W1):W199–W205.

Lis, R., Karrasch, C. C., Poulos, M. G., Kunar, B., Redmond, D., Duran, J. G., Badwe, C. R., Schachterle, W., Ginsberg, M., Xiang, J., et al. (2017). Conversion of adult endothelium to immunocompetent haematopoietic stem cells. Nature, 545(7655):439–445.

McClard, C. K., Kochukov, M. Y., Herman, I., Liu, Z., Eblimit, A., Moayedi, Y., Ortiz-Guzman, J., Colchado, D., Pekarek, B., Panneerselvam, S., et al. (2018). POU6f1 mediates neuropeptide-dependent plasticity in the adult brain. Journal of Neuroscience, 38(6):1443–1461.

Melé, M., Ferreira, P. G., Reverter, F., DeLuca, D. S., Monlong, J., Sammeth, M., Young, T. R., Goldmann, J. M., Pervouchine, D. D., Sullivan, T. J., et al. (2015). The human transcriptome across tissues and individuals. Science, 348(6235):660–665.

Ott, C. J., Blackledge, N. P., Kerschner, J. L., Leir, S. H., Crawford, G. E., Cotton, C. U., and Harris, A. (2009). Intronic enhancers coordinate epithelial-specific looping of the active CFTR locus. Proceedings of the National Academy of Sciences of the United States of America, 106(47):19934–19939.

Pennacchio, L. A., Loots, G. G., Nobrega, M. A., and Ovcharenko, I. (2007). Predicting tissue-specific enhancers in the human genome. Genome Research, 17(2):201–211.

Pervouchine, D. D., Djebali, S., Breschi, A., Davis, C. A., Barja, P. P., Dobin, A., Tanzer, A., Lagarde, J., Zaleski, C., See, L. H., et al. (2015). Enhanced transcriptome maps from multiple mouse tissues reveal evolutionary constraint in gene expression. Nature Communications, 6(1):1–11.

Quinlan, A. R. and Hall, I. M. (2010). BEDTools: a flexible suite of utilities for comparing genomic features. Bioinformatics, 26(6):841–842.

Rand, E. and Cedar, H. (2003). Regulation of imprinting: A multi-tiered process. Journal of Cellular Biochemistry, 88(2):400–407.

Rose, A. B. (2019). Introns as Gene Regulators: A Brick on the Accelerator. Frontiers in Genetics, 9(FEB):672.

Rué, P. and Martinez Arias, A. (2015). Cell dynamics and gene expression control in tissue homeostasis and development. Molecular Systems Biology, 11(2):792.

Schmitt, A. D., Hu, M., Jung, I., Xu, Z., Qiu, Y., Tan, C. L., Li, Y., Lin, S., Lin, Y., Barr, C. L., et al. (2016). A Compendium of Chromatin Contact Maps Reveals Spatially Active Regions in the Human Genome. Cell Reports, 17(8):2042–2059.

Shaul, O. (2017). How introns enhance gene expression.

Shlyueva, D., Stampfel, G., and Stark, A. (2014). Transcriptional enhancers: From properties to genome-wide predictions.

Vuong, C. K., Black, D. L., and Zheng, S. (2016). The neurogenetics of alternative splicing.

Zabidi, M. A., Arnold, C. D., Schernhuber, K., Pagani, M., Rath, M., Frank, O., and Stark, A. (2015). Enhancer-core-promoter specificity separates developmental and housekeeping gene regulation. Nature, 518(7540):556–559.

